# Distinct complexes of RAD51 paralogs participate in different fork protection pathways

**DOI:** 10.64898/2026.04.29.721569

**Authors:** Debanjali Bhattacharya, Harsh Kumar Dwivedi, Satyaranjan Sahoo, Shariva Kadupatil, Ganesh Nagaraju

## Abstract

Error-free genome duplication is critical for cellular homeostasis and genome maintenance. Stalled forks undergo remodeling during replication stress, and unprotected forks undergo degradation by nucleases. RAD51 paralogs are evolutionarily conserved essential proteins for genome maintenance with diverse roles ranging from homologous recombination (HR) to replication stress responses. However, the mechanisms underlying their participation in fork maintenance remain unclear. Here, we demonstrate that the RAD51 paralogs do not participate in the canonical SMARCAL1-BRCA2 axis of fork protection. Instead, RAD51D–XRCC2 (DX2) and RAD51C–XRCC3 (CX3) complexes protect forks remodeled by the FBH1 helicase, whereas the RAD51B–RAD51C (BC) subcomplex but not BRCA2 safeguards forks remodeled by the FANCM translocase, revealing a new FANCM-mediated pathway of fork remodeling which is protected by the BC sub-complex. Mechanistically, we show that FANCM-mediated fork reversal is RAD51-dependent and generates substrates for MRE11-, EXO1-, and DNA2-mediated degradation in the absence of RAD51B/C. Our findings establish the participation of the RAD51 paralogs in multiple fork protection pathways in a fork-remodeler-specific manner, highlighting the existence of several independent fork remodeling and protection mechanisms for maintaining genomic stability under replication stress.

## Introduction

Accurate and timely completion of genome duplication during the cell cycle is essential for genome maintenance. However, the replicating forks can encounter various endogenous and exogenous challenges, including DNA lesions, limiting nucleotide pools, repetitive DNA regions, DNA secondary structures, transcription-replication conflicts (TRCs), and DNA-protein crosslinks, leading to fork stalling and replication stress (Cortez, 2019; Nagaraju & Scully, 2007; Zeman & Cimprich, 2014). Cells have evolved with several defense mechanisms to safeguard the genome during replication stress. Fork reversal is a prominent cellular mechanism for dealing with impediments to the ongoing replication machinery. Stalled replication forks undergo fork reversal, generating a four-way junction, which slows down DNA replication, stabilizes the stalled forks and facilitates the repair of template lesions (Berti *et al*, 2020a; Neelsen & Lopes, 2015; Quinet *et al*, 2017). Further, reversed forks also promote fork restart through template switching or by MUS81-dependent mechanisms (Cortez, 2015; Hanada *et al*, 2007; Pasero & Vindigni, 2017). Although reversed forks are beneficial for overcoming replication stress, they are susceptible to degradation by several nucleases and hence require protection by various proteins. In the absence of proper fork protection mechanisms, fork reversal can become a source of replication stress (Rickman & Smogorzewska, 2019; Saxena & Zou, 2022).

Multiple mechanistically distinct pathways of replication fork remodeling have been reported. The most extensively characterized pathway involves the SNF2-family DNA translocases, including SMARCAL1, ZRANB3, and HLTF, which act in a common pathway of fork reversal. Forks reversed by the SNF2-remodelers are subsequently protected by BRCA1, BRCA2, and components of the Fanconi anemia (FA) pathway such as FANCD2 (Liao *et al*, 2018; Michl *et al*, 2016; Schlacher *et al*, 2011; Schlacher *et al*, 2012; Taglialatela *et al*, 2017). In this context, BRCA2-mediated stabilization of RAD51 nucleofilaments is critical to prevent pathological degradation of nascent DNA by nucleases such as MRE11 and EXO1. In addition, an alternative pathway of fork remodeling has been described in which the FBH1 helicase promotes fork reversal independently of SNF2-family remodelers, where fork protection is mediated by factors including 53BP1 and BOD1L rather than BRCA2 (Liu *et al*, 2020). Notably, FBH1-remodeled forks exhibit distinct nuclease sensitivities, being preferentially degraded by DNA2, highlighting fundamental differences in the processing of reversed fork structures generated by different remodelers. Emerging evidence also suggests the existence of additional fork remodeling factors, including PICH, which has been shown to mediate extensive fork remodeling downstream of the SNF2-fork remodelers (Tian *et al*, 2021). Additionally, biochemical studies have demonstrated the fork remodeling activity of several DNA translocases/helicases in vitro; including RAD54, BLM-WRN complex and FANCM (Bugreev *et al*, 2011; Gari *et al*, 2008; Machwe *et al*, 2006), although their roles in fork reversal in vivo remain obscure. Collectively, these studies support the existence of multiple, parallel pathways of fork remodeling and protection in cells, although the molecular mechanism and genomic context of pathway choice remain unknown.

Many DNA damage repair proteins, including BRCA1/2, RAD51, RAD51 paralogs, 53BP1 and FA pathway components contribute to the protection of stalled replication forks (Rickman & Smogorzewska, 2019; Tye *et al*, 2021). While the roles of RAD51 and BRCA proteins in fork protection have been extensively characterized (Hashimoto *et al*, 2010; Schlacher *et al*., 2011; Schlacher *et al*., 2012), the mechanisms underlying RAD51 paralogs participation in replication stress responses remain incompletely understood. In mammalian cells, five RAD51 paralogs—RAD51B, RAD51C, RAD51D, XRCC2, and XRCC3—assemble into distinct complexes, most notably the BCDX2, CX3 and the BC and DX2 subcomplexes (Dosanjh *et al*, 1998; Greenhough *et al*, 2023; Kurumizaka *et al*, 2002; Lio *et al*, 2003; Liu *et al*, 2002; Longo *et al*, 2023; Masson *et al*, 2001a; Masson *et al*, 2001b; Miller *et al*, 2002; Rawal *et al*, 2023; Sigurdsson *et al*, 2001). The RAD51 paralogs are best known for their canonical function as mediators of RAD51 nucleofilament formation during HR (Chun *et al*, 2013; Liu *et al*, 2007; Masson *et al*., 2001b; Nagaraju *et al*, 2009; Nagaraju *et al*, 2006; Sigurdsson *et al*., 2001; Takata *et al*, 2000; Takata *et al*, 2001; Taylor *et al*, 2015). However, emerging evidence has demonstrated the involvement of RAD51 paralogs in a range of replication-associated processes, including intra-S-phase checkpoint signaling, replication fork progression during dNTP pool fluctuations, fork protection and its restart, and resolution of R-loops (Bhattacharya *et al*, 2022; Sahoo *et al*, 2026; Saxena *et al*, 2019; Saxena *et al*, 2018; Somyajit *et al*, 2013; Somyajit *et al*, 2015; Somyajit *et al*, 2012). Loss of individual RAD51 paralogs leads to embryonic lethality in mice, and mutations of the paralogs are associated with breast and ovarian cancers, and FA/FA-like disorder, highlighting the importance of these proteins in genome maintenance (Deans *et al*, 2000; Kuznetsov *et al*, 2009; Loveday *et al*, 2011; Meindl *et al*, 2010; Pittman & Schimenti, 2000; Prakash *et al*, 2021; Shu *et al*, 1999; Somyajit *et al*, 2010; Vaz *et al*, 2010). Earlier studies in mammalian cells demonstrated that loss of RAD51C, XRCC2, or XRCC3 leads to defects in stalled fork protection, implicating both the BCDX2 and CX3 complexes in the fork stabilization (Bhattacharya *et al*, 2025; Somyajit *et al*., 2015). In addition, pathological mutants of RAD51C have been reported to exhibit fork protection defect (Kolinjivadi *et al*, 2023; Longo *et al*., 2023; Somyajit *et al*., 2015). Despite these advances, the molecular mechanism by which the different RAD51 paralog complexes participate in fork protection remains largely unknown.

Here, we report the participation of distinct complexes of RAD51 paralogs in protecting stalled replication forks. We find that, in contrast to BRCA2-deficient cells, fork degradation in RAD51 paralog-deficient cells cannot be rescued by co-depletion of SMARCAL1, indicating that the RAD51 paralogs participate in fork protection independently of the canonical SNF2-mediated pathway. Notably, we find that DX2 and CX3 complexes protect forks remodelled by the FBH1 helicase, whereas BC sub-complex safeguard forks remodelled by the FANCM DNA translocase, revealing a previously uncharacterized pathway of fork remodelling and protection. Mechanistically, we show that FANCM-mediated fork reversal is RAD51-dependent and undergoes fork degradation by the MRE11, DNA2 and EXO1 nucleases in the absence of RAD51B/C-mediated fork protection. Finally, we demonstrate that FANCM and RAD51B/C-deficient cells accumulate replication stress-associated DNA damage, and co-depletion of FANCM in RAD51B/C-depleted cells rescues the DNA damage levels but not cell survival, highlighting the essential functions of these proteins during replication stress response. Overall, our findings identify that distinct RAD51 paralog complexes protect stalled forks in a remodeler-specific manner to maintain genome stability during replication stress.

## Results

### BRCA2 but not the RAD51 paralogs protect forks remodeled by the SNF2-family remodelers

The role of the RAD51 paralogs in stalled fork protection was reported about a decade ago (Somyajit *et al*., 2015). Pathological variants of the RAD51 paralogs, when expressed in cells, have been shown to cause fork degradation (Kolinjivadi *et al*., 2023; Longo *et al*., 2023; Somyajit *et al*., 2015). Nevertheless, a mechanistic analysis of the role of all five RAD51 paralogs in replication fork protection has not been done to date. To examine the role of the RAD51 paralogs in stalled fork protection, we depleted each of RAD51B, RAD51C, RAD51D, XRCC2 and XRCC3 in U2OS and RPE1 cells using shRNA-mediated gene knockdown (Figure S1A). The paralog-depleted cells were sequentially pulse-labelled with the thymidine analogs CldU and IdU, subjected to prolonged replication stress by HU treatment and processed for the DNA fiber assay to study fork protection. RAD51B-, RAD51C-, RAD51D-, XRCC2, and XRCC3-depleted cells showed a reduced IdU/CldU ratio as compared to control cells, indicating that in the absence of the RAD51 paralogs, stalled forks undergo extensive fork degradation in both U2OS and RPE1 cells (Figures 1A-1C). Fork degradation occurs downstream to fork reversal when fork protection factors are compromised. BRCA2 is known to protect forks reversed by the SNF2 family remodelers, namely SMARCAL1, ZRANB3 and HLTF, all of which participate in a common pathway of fork reversal (Schlacher *et al*., 2011; Taglialatela *et al*., 2017). We questioned whether the RAD51 paralogs also participate in the protection of stalled forks remodeled by SNF2 family remodelers. To test this, we co-depleted SMARCAL1 with each RAD51 paralog in U2OS cells and performed a fork protection assay. Surprisingly, co-depletion of RAD51 paralogs with SMARCAL1 failed to rescue fork degradation in any of the paralog-depleted cells, as indicated by the reduced IdU to CldU ratios (Figure 1D and 1E). In contrast, SMARCAL1-depletion completely rescued fork degradation occurring in BRCA2-deficient cells (Figure 1D, 1E and S1B), indicating that BRCA2, but not the RAD51 paralogs, participates in the protection of stalled forks remodeled by SMARCAL1. Additionally, fork degradation in RAD51C-deficient cells could not be rescued by co-depletion of either SMARCAL1, ZRANB3, or HLTF (Figure S1C-S1G). These results indicate that the RAD51 paralogs are essential for fork protection during replication stress; however, they do not participate in the SNF2-BRCA2 axis of fork remodeling/protection.

**Figure 1.**
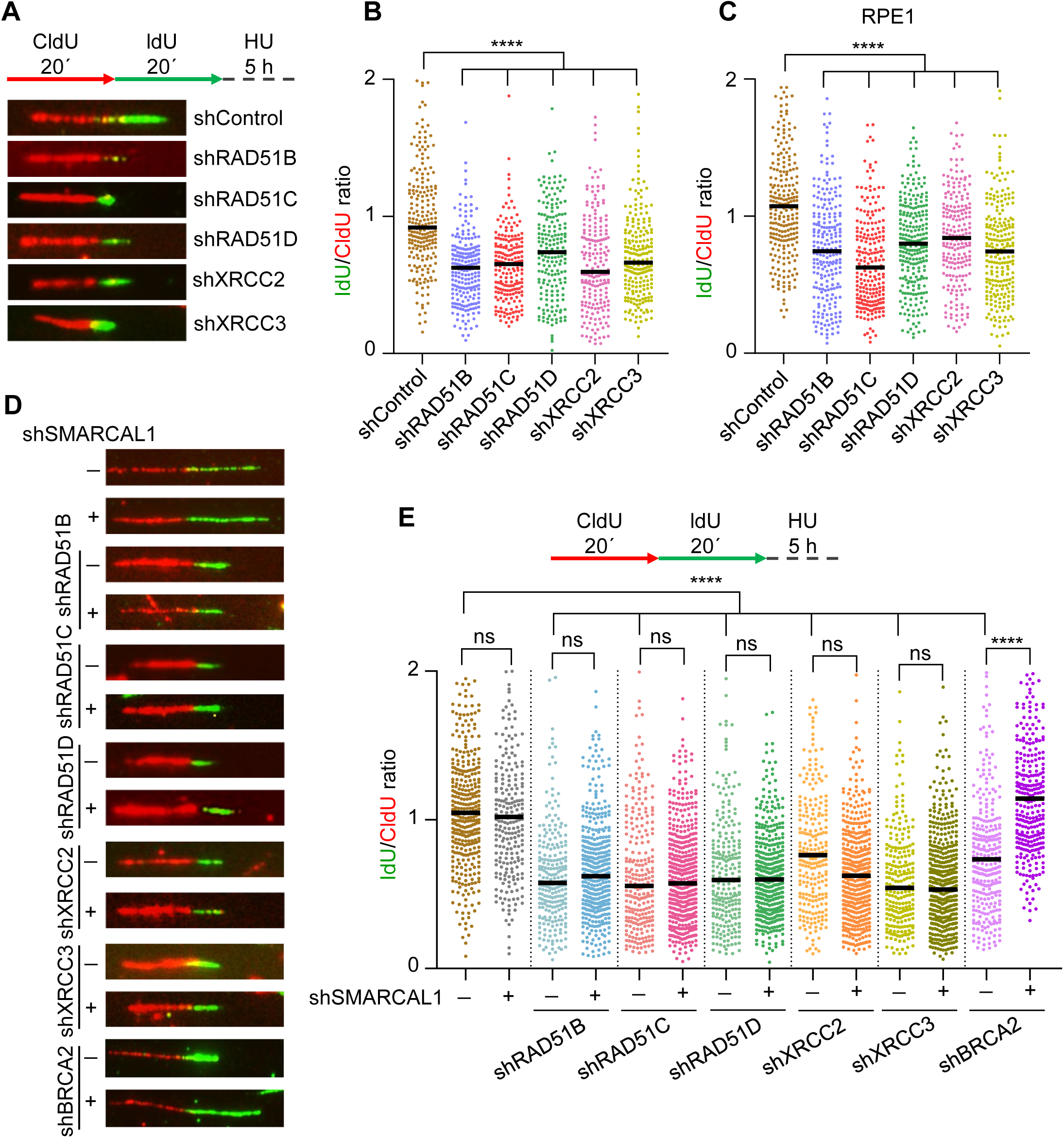
The RAD51 paralogs do not participate in the SNF2-BRCA2 axis of fork protection. (A) Schematic of pulse-labeling (top) and representative DNA fiber images showing fork degradation in control and RAD51 paralog-depleted U2OS cells (bottom). (B) Quantitative scatter plot of IdU/CldU track length ratios in U2OS cells as indicated in (A). Black bars represent median values. Number of fibers (n) calculated from three independent experiments are: shControl-226; shRAD51B-220; shRAD51C-201; shRAD51D-196; shXRCC2-202; shXRCC3-216. Mann-Whitney *t* test, ****P<0.0001, P(shControl vs shRAD51B; shControl vs shRAD51C; shControl vs shRAD51D; shControl vs shXRCC2; shControl vs shXRCC3) <0.0001. (C) Scatter plot showing IdU/CldU ratios in control vs RAD51 paralog-depleted RPE1 cells treated with 4 mM HU for 5 h. n calculated from three independent experiments are: shControl-240; shRAD51B-235; shRAD51C-228; shRAD51D-245; shXRCC2-211; shXRCC3-234. Black bars represent median values. Mann-Whitney *t* test, ****P<0.0001, P(shControl vs shRAD51B; shControl vs shRAD51C; shControl vs shRAD51D; shControl vs shXRCC2; shControl vs shXRCC3) <0.0001. (D) Representative DNA fibers of fork degradation assay in U2OS cells treated with the indicated shRNAs. (E) Schematic of experimental design (top) and quantitative scatter plot showing IdU/CldU ratios in cells as shown in (D). Black bars represent median values. n calculated from three biological replicates are: shControl-356; shSMARCAL1-242; shRAD51B-245; shRAD51B+shSMARCAL1-427; shRAD51C-250; shRAD51C+shSMARCAL1-467; shRAD51D-233; shRAD51D+shSMARCAL1-424; shXRCC2-248; shXRCC2+shSMARCAL1-431; shXRCC3-233; shXRCC3+shSMARCAL1-478; shBRCA2- 311; shBRCA2+shSMARCAL1-313. Mann-Whitney *t* test, ****P<0.0001, ns- non-significant, P(shControl vs shRAD51B; shControl vs shRAD51C; shControl vs shRAD51D; shControl vs shXRCC2; shControl vs shXRCC3; shBRCA2 vs shBRCA2+shSMARCAL1) <0.0001.

### The DX2 and CX3 paralog complexes protect forks remodeled by the FBH1 helicase

A recent study found that the FBH1 helicase remodels stalled replication forks independently of the SNF2-remodelers pathway, and these forks are protected by proteins such as 53BP1 and BOD1L (Liu *et al*., 2020). Since our results showed that the RAD51 paralogs do not participate in the SMARCAL1-BRCA2 axis, we tested whether the RAD51 paralogs participate in the protection of stalled forks remodeled by the FBH1 helicase. To this end, we co-depleted each of the five RAD51 paralogs with FBH1 and assessed their effect on fork degradation by the DNA fiber assay. Surprisingly, FBH1 co-depletion rescued fork degradation in RAD51D, XRCCC2, and XRCC3-depleted cells (Figures 2A and 2B). RAD51C-depleted cells showed a partial but significant rescue of fork degradation upon FBH1 co-depletion (Figures 2A, 2B, S1H). Consistent with previous studies, fork degradation in 53BP1- and BOD1L-depleted cells were rescued by co-depletion of FBH1 (Figures 2A, 2B, S1I). These results point to a functional separation between the different RAD51 paralog complexes and establish that the RAD51D-XRCC2 subcomplex, as well as the RAD51C-XRCC3 complex, or alternatively the X3CDX2 complex, protects forks remodeled by the FBH1 helicase.

**Figure 2.**
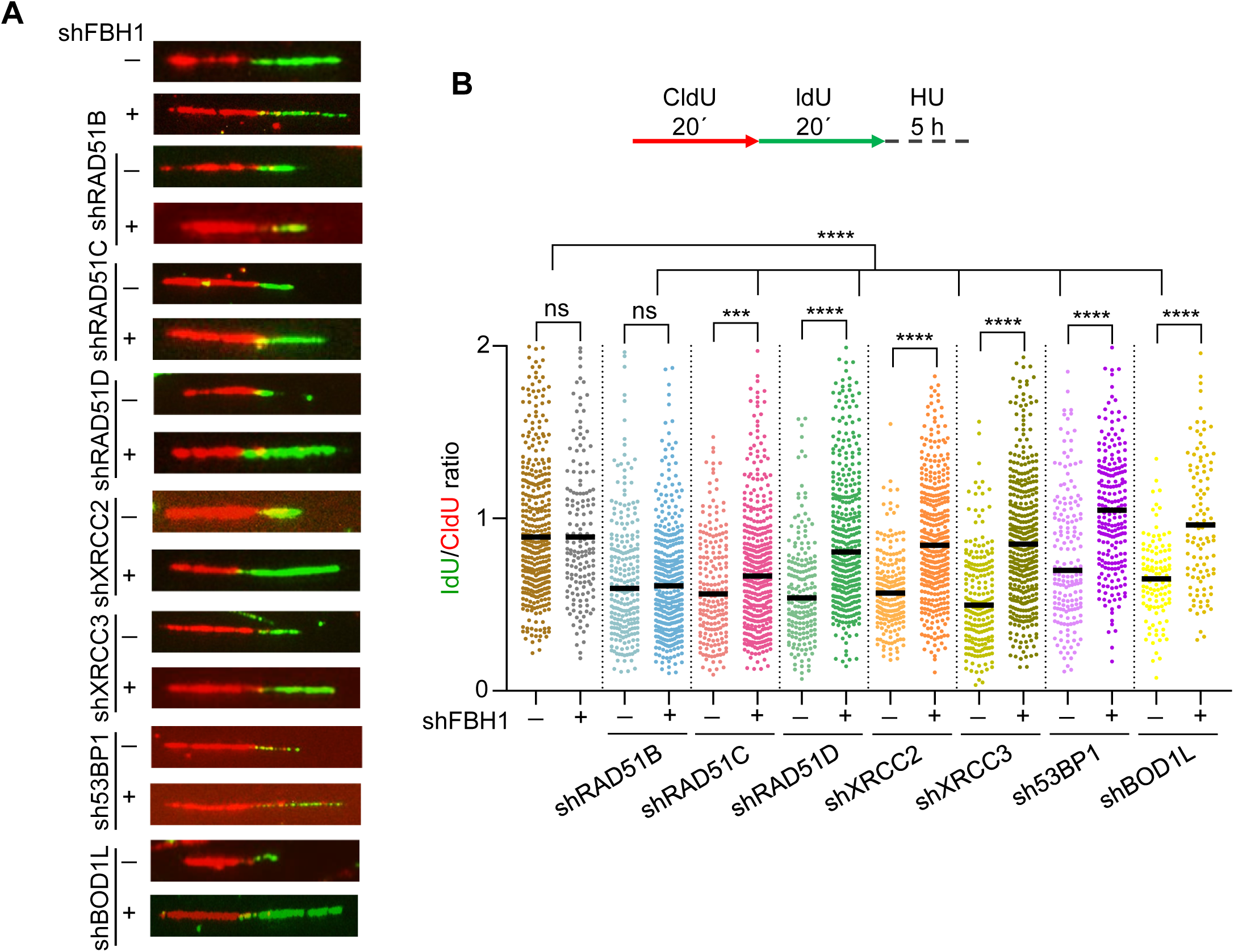
The DX2 and CX3 complexes protect FBH1 helicase remodeled forks. (A) Representative DNA fiber images in U2OS cells showing fork degradation in the indicated samples. (B) Schematic of pulse-labeling scheme (top) and scatter plot of IdU/CldU track lengths in samples as indicated in (A) (bottom). Black bars represent median values. n from three independent experimental repeats are: shControl-404; shFBH1-173; shRAD51B-233; shRAD51B+shFBH1-380; shRAD51C-197; shRAD51C+shFBH1-398; shRAD51D-192; shRAD51D+shFBH1-386; shXRCC2-202; shXRCC2+shFBH1-420; shXRCC3-206; shXRCC3+shFBH1-398; sh53BP1-180; sh53BP1+shFBH1-227; shBOD1L-111; shBOD1L+shFBH1- 114. Mann-Whitney *t* test, ****P<0.0001, ***P<0.001, ns- non-significant, P(shControl vs shRAD51B; shControl vs shRAD51C; shControl vs shRAD51D; shControl vs shXRCC2; shControl vs shXRCC3; shRAD51D vs shRAD51D+shFBH1; shXRCC2 vs shXRCC2+shFBH1; shXRCC3 vs shXRCC3+shFBH1; sh53BP1 vs sh53BP1+shFBH1; shBOD1L vs shBOD1L+shFBH1) <0.0001, P(shRAD51C vs shRAD51C+shFBH1) = 0.0010.

### RAD51B-RAD51C subcomplex protects stalled forks remodeled by the FANCM DNA translocase

The failure of SMARCAL1 or FBH1 co-depletion to rescue fork degradation in RAD51B-deficient cells, as well as the partial rescue of fork degradation in RAD51C-deficient cells with FBH1 co-depletion, indicated the likely existence of a third, independent pathway of fork reversal and protection. FANCM is a large scaffolding protein with DNA translocase activity and has been shown to mediate fork reversal in vitro (Gari *et al*., 2008). We thus hypothesized that the RAD51 paralogs may protect stalled forks remodeled by FANCM DNA translocase. To test this, we performed the fork protection assay in RAD51 paralogs and FANCM co-depleted U2OS cells. Interestingly, fork degradation in RAD51B- and RAD51C-depleted cells was rescued upon FANCM co-depletion, but not in RAD51D-, XRCC2-, or XRCC3-depleted cells (Figures 3A, 3B, S2A-S2E). The same result was obtained in RPE1 cells, where FANCM knockdown rescued fork degradation in RAD51B and RAD51C-deficient cells, but not in RAD51D, XRCC2 or XRCC3-deficient cells (Figures S2F and S2G). We independently assessed fork degradation in RAD51C+FANCM and RAD51C+SMARCAL1-depleted RPE1 cells. In accordance with our previous observations in U2OS cells (Figures 1D, 1E, 3A and 3B), the fork protection defect in RAD51C-deficient cells was rescued by FANCM but not SMARCAL1 knockdown (Figure S2H and S2I). To further confirm our results, we tested whether FANCM depletion can rescue fork degradation in RAD51C and SMARCAL1/ZRANB3/HLTF-co-depleted cells. As observed earlier (Figures S1F and S1G), depletion of SMARCAL1/ZRANB3/HLTF failed to rescue fork degradation in RAD51C-deficient cells; however, depletion of FANCM in these cells rescued the fork protection phenotype (Figures S3A and S3B). A similar observation was made in RAD51C and FBH1-depleted cells, in which co-depletion of FANCM completely rescued the fork degradation observed in RAD51C or RAD51C+FBH1-depleted cells (S3C and S3D). These results indicate that the RAD51B-RAD51C subcomplex participates in a novel pathway of fork protection and protects forks remodeled by the FANCM DNA translocase. Next, we tested whether the FANCM-mediated pathway is epistatic to the previously defined pathways of fork remodeling. To test this, we performed fork protection assays by FANCM knockdown in BRCA2- or 53BP1/BOD1L-depleted cells and measured the fork stability. Surprisingly, FANCM co-depletion failed to rescue the fork degradation in either BRCA2, 53BP1, or BOD1L-depleted U2OS cells (Figures 3C, 3D, S3E and S3F), indicating that FANCM functions independently of the SNF2 or FBH1-mediated remodeling pathway. 53BP1 was previously shown to be dispensable for fork protection in hTERT-RPE1 cells (Liu *et al*., 2020). In our experiments with RPE1 cells, FANCM co-depletion completely rescued the fork degradation occurring in RAD51C-depleted cells but not in BOD1L-depleted cells (Figures 3E and 3F). However, in contrast to U2OS cells, FANCM co-depletion partially rescued the fork degradation in BRCA2-deficient RPE1 cells (Figures 3E and 3F), indicating cell-line-specific effects of FANCM depletion in a BRCA2-deficient background. As our observations indicated that the BC sub-complex participates in stalled fork protection independently of the BRCA2- or 53BP1-mediated pathways, we expected an additive effect of fork degradation in the paralog and BRCA2 or 53BP1 co-depleted cells. However, our analysis in U2OS cells revealed a similar level of fork degradation in RAD51C/BRCA2, as well as RAD51C/53BP1 single- and double-depleted cells (Figures S4A-S4F). These results corroborate a previous study showing that co-depletion of 53BP1 and BRCA2 failed to alter fork degradation levels, possibly because the assay was unable to detect additive changes in fork degradation (Liu *et al*., 2020). Taken together, our studies indicate that FANCM, along with RAD51B-RAD51C, participates in a novel pathway of fork remodeling/protection, which is independent of both the BRCA2- and 53BP1/BOD1L-dependent pathways.

**Figure 3.**
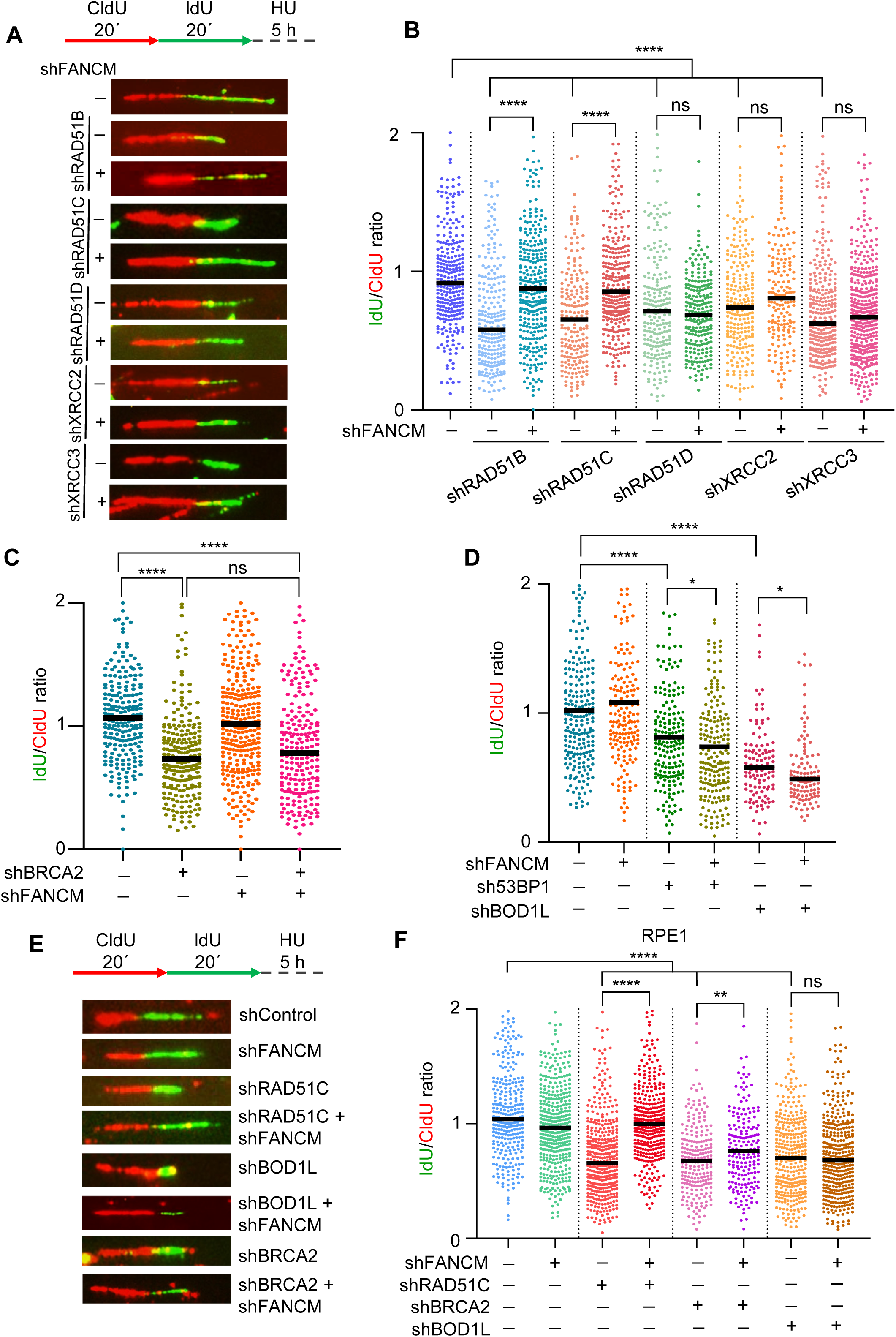
BC subcomplex participates in a novel pathway of fork protection. (A) DNA fiber assay schematic (top) and representative images showing fork degradation in Control, RAD51 paralogs- and RAD51 paralogs + FANCM-depleted U2OS cells. (B) Scatter plot showing quantification of IdU to CldU ratios in cells as indicated in (A). Black bars represent median values. n from three independent experimental repeats are: shControl-286; shRAD51B-247; shRAD51B+shFANCM-336; shRAD51C- 214; shRAD51C+shFANCM-305; shRAD51D-223; shRAD51D+shFANCM-236; shXRCC2-234; shXRCC2+shFANCM-178; shXRCC3-223; shXRCC3+shFANCM-422. Mann-Whitney *t* test, ****P<0.0001, ns- non-significant, P(shControl vs shRAD51B; shControl vs shRAD51C; shControl vs shRAD51D; shControl vs shXRCC2; shControl vs shXRCC3; shRAD51B vs shRAD51B+shFANCM; shRAD51C vs shRAD51C+shFANCM) <0.0001. (C) Scatter plot of IdU/CldU ratios from DNA fiber assay in Control, BRCA2, FANCM and BRCA2+FANCM-depleted U2OS cells. Black bars represent median values. n from three independent experimental repeats are as follows: shControl-228; shBRCA2-237; shFANCM-312; shBRCA2+shFANCM-219. Mann-Whitney *t* test, ****P<0.0001, ns- non-significant, P(shControl vs shBRCA2; shControl vs shBRCA2+shFANCM) <0.0001. (D) Scatter plot of IdU/CldU ratios from DNA fiber assay in Control, FANCM, 53BP1, BOD1L, 53BP1+FANCM and BOD1L+FANCM-depleted U2OS cells. Black bars represent median values. n counted from three independent experimental repeats are: shControl-221; shFANCM-179; sh53BP1-189; sh53BP1+shFANCM-198; shBOD1L-112; shBOD1L+shFANCM-117. Mann-Whitney *t* test, ****P<0.0001, *P<0.05, ns-non-significant, P(shControl vs sh53BP1; shControl vs shBOD1L) <0.0001, (sh53BP1 vs sh53BP1+shFANCM) = 0.0327, (shBOD1L vs shBOD1L+shFANCM) = 0.0158. (E) Pulse-labeling schematic (top) and representative DNA fibers showing fork degradation in the indicated samples in RPE1 cells. (F) Scatter plot of IdU/CldU ratios in RPE1 cells as indicated in (E). Black bars represent median values. n counted from three independent experiments are as follows: shControl-310; shFANCM-423; shRAD51C-391; shRAD51C+shFANCM-354; shBRCA2-224; shBRCA2+shFANCM-196; shBOD1L-359; shBOD1L+shFANCM-376. Mann-Whitney *t* test, ****P<0.0001, **P<0.01, ns- non-significant, P(shControl vs shRAD51C; shControl vs shBRCA2; shControl vs shBOD1L; shRAD51C vs shRAD51C+shFANCM) <0.0001, P(shBRCA2 vs shBRCA2+shFANCM) = 0.0073.

### FANCM-mediated fork reversal is dependent on RAD51 and undergoes MRE11, DNA2 and EXO1-mediated degradation

Previous studies have shown that RAD51 promotes fork reversal under genotoxic stress and participates upstream of the fork remodelers to initiate fork reversal via its strand-exchange activity (Liu *et al*, 2023; Zellweger *et al*, 2015). To test whether RAD51 is essential for fork reversal in the FANCM-mediated pathway, we depleted RAD51 along with RAD51B and RAD51C and examined the fork degradation occurring in these cells. RAD51-depletion rescued fork degradation in RAD51B and RAD51C-deficient cells (Figure 4A, S4G and S4H), suggesting that fork remodeling by FANCM is RAD51-dependent. PARP protein promotes fork reversal by inhibiting RECQL1’s fork restart activity (Berti *et al*, 2013; Ray Chaudhuri *et al*, 2012; Zellweger *et al*., 2015). Inhibiting PARP with olaparib treatment significantly rescued the fork protection defect in RAD51B and RAD51C-depleted cells (Figure 4B), further confirming that fork reversal leads to fork degradation in these cells. Since fork protection is dependent on stable RAD51 nucleofilament formation, overexpression of wild-type RAD51 (WT) or II3A RAD51 mutant, which is competent for stable nucleofilament formation but deficient in strand exchange activity, rescued fork degradation in RAD51B and RAD51C- deficient cells. In contrast, T131P RAD51, which cannot form stable nucleofilaments, failed to rescue fork degradation (Figure 4C, S4I and S4J).

**Figure 4.**
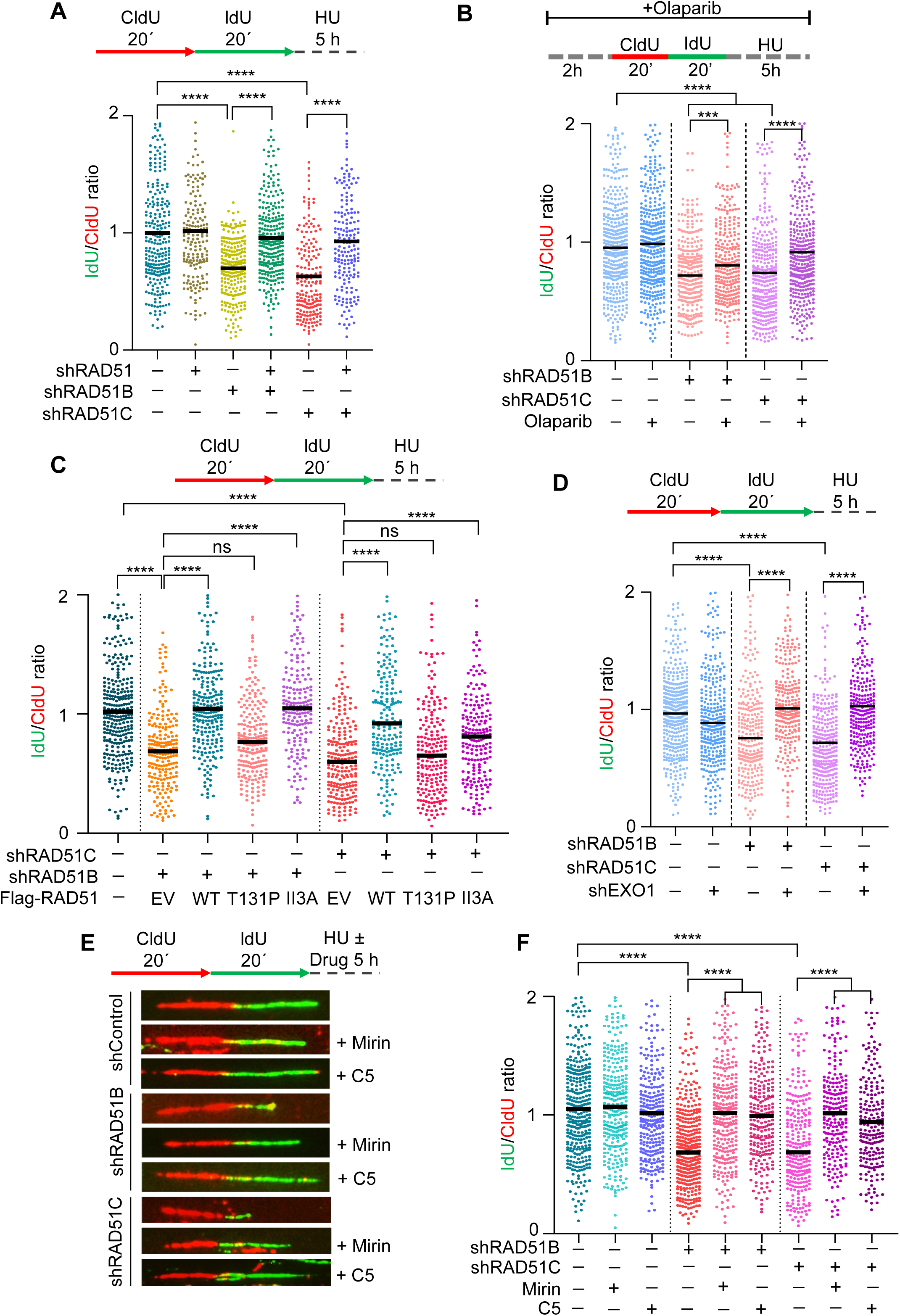
Fork remodeling by FANCM is RAD51-dependent and susceptible to degradation by multiple nucleases. (A) Schematic of pulse-labeling and scatter plot depicting IdU/CldU ratios from the indicated samples in U2OS cells. Black bars represent mean values. The total number of fibers counted from three independent biological replicates are: shControl-264; shRAD51B-231; shRAD51C-185; shRAD51-181; shRAD51B+shRAD51-275; shRAD51C+shRAD51-185. Mann-Whitney *t* test, ****P<0.0001, ns- non-significant, P(shControl vs shRAD51C; shControl vs shRAD51B; shRAD51B vs shRAD51B+shRAD51; shRAD51C vs shRAD51C+shRAD51) <0.0001. (B) Schematic of pulse-labeling and olaparib treatment and scatter plot depicting IdU/CldU ratios from the indicated samples in U2OS cells. Black bars represent median values. The total number of fibers counted from three independent biological replicates are: shControl-378; shControl+olaparib-348; shRAD51B-203; shRAD51B+olaparib-179; shRAD51C-335; shRAD51C+olaparib-369. Mann-Whitney *t* test, ****P<0.0001, ns- non-significant, P(shControl vs shRAD51B; shControl vs shRAD51C; shRAD51C vs shRAD51C+olaparib) <0.0001, P(shRAD51B vs shRAD51B+olaparib) = 0.0002. (C) Schematic of pulse-labeling and scatter plot depicting IdU/CldU ratios from the indicated samples in U2OS cells. Black bars represent median values. EV- empty vector, WT- wild type. The total number of fibers counted from three independent biological replicates are: shControl-301; shRAD51B+EV-213; shRAD51B+WT 244; shRAD51B+ T131P-218; shRAD51B+ II3A RAD51-193; shRAD51C+EV-214; shRAD51C+WT RAD51-199; ; shRAD51C+ T131P-207; shRAD51C+ II3A RAD51-206. Mann-Whitney *t* test, ****P<0.0001, ns- non-significant, P(shControl vs shRAD51C; shControl vs shRAD51B; shRAD51B+EV vs shRAD51B+WT RAD51; shRAD51B+EV vs shRAD51B+II3A RAD51; shRAD51C+EV vs shRAD51C+WT RAD51; shRAD51C+EV vs shRAD51C+II3A RAD51) <0.0001. (D) Schematic of pulse-labeling and scatter plot depicting IdU/CldU ratios from fork protection assay in U2OS cells treated with the indicated shRNAs. Black bars represent median values. The total number of fibers counted from three independent biological replicates are: shControl-237; shEXO1-315; shRAD51B-277; shRAD51B+shEXO1-268; shRAD51C-239; shRAD51C+shEXO1-263. Mann-Whitney *t* test, ****P<0.0001, ns- non-significant, P(shControl vs shRAD51B; shControl vs shRAD51C; shRAD51B vs shRAD51B+shEXO1; shRAD51C vs shRAD51C+shEXO1) <0.0001. (E) Pulse labeling schematic and representative DNA fiber images showing fork protection in control, RAD51B and RAD51C-depleted cells treated with mirin or C5 inhibitors. (F) Scatter plot representing IdU/CldU ratios in cells as mentioned in (E). The black bars represent median values. The total number of fibers counted from three independent biological replicates are: shControl-264; shRAD51B-368; shRAD51B+Mirin303;- shRAD51B+C5-247; shRAD51C-259; shRAD51C+Mirin-227; shRAD51C+C5-222. Mann-Whitney *t* test, ****P<0.0001, ns- non-significant, P(shControl vs shRAD51B; shControl vs shRAD51C; shRAD51B vs shRAD51B+Mirin; shRAD51B vs shRAD51B+C5; shRAD51C vs shRAD51C+Mirin; shRAD51C vs shRAD51C+C5) <0.0001.

Nucleases such as MRE11, EXO1, and DNA2 have been shown to degrade reversed fork structures (Kolinjivadi *et al*, 2017a; Lemacon *et al*, 2017; Taglialatela *et al*., 2017; Thangavel *et al*, 2015). Depletion of the EXO1 endonuclease inhibited fork degradation in RAD51B/C-deficient cells (Figure 4D, S4K and S4L). Parallelly, inhibition of MRE11 and DNA2 nucleases using mirin and C5 inhibitors, respectively, prevented fork degradation in both RAD51B and RAD51C-deficient cells (Figures 4E and 4F), indicating that forks reversed by FANCM are susceptible to MRE11, DNA2 and EXO1-mediated degradation when fork protection factors are absent. Together, our results indicate that FANCM-mediated fork reversal is RAD51-dependent and the BC sub-complex protects such forks from the action of multiple nucleases.

### FANCM-mediated fork remodeling leads to DNA damage in RAD51B and RAD51C-deficient cells

In the absence of fork protection, reversed forks can collapse into double-strand breaks (DSBs), and fork degradation can result in the accumulation of single-stranded DNA (ssDNA) (Bhattacharya *et al*., 2025; Kolinjivadi *et al*., 2017a; Schlacher *et al*., 2011; Taglialatela *et al*., 2017). To test whether FANCM-mediated fork reversal leads to the accumulation of DNA damage in RAD51B and RAD51C-deficient cells under replication stress, we treated cells with HU to induce replication stress and performed immunofluorescence experiments, scoring γH2AX and RPA70 foci as markers of DSBs and ssDNA, respectively. RAD51B-deficient cells showed a significant increase in γH2AX and RPA70 foci, however, RAD51C-depleted cells accumulated the highest amount DNA damage foci upon HU treatment, which were significantly rescued by FANCM-co-depletion (Figures 5A-5C), demonstrating that FANCM- mediated fork reversal leads to the accumulation of DNA damage in the absence of fork protection. The higher levels of DNA damage in RAD51C-depleted cells possibly reflect additional functions of RAD51C during replication stress, such as R-loop resolution (Sahoo *et al*., 2026) or fork protection in the FBH1-mediated pathway. However, survival assays with RAD51B/C and FANCM-depleted cells revealed that the cell survival defect in RAD51B or RAD51C-depleted cells was comparable to RAD51B/RAD51C+FANCM co-depleted cells under continuous HU and APH treatment (Figures 5D and 5E). Interestingly, consistent with our previous results (Figure 5A-5C), RAD51C-deficient cells exhibited a marginally enhanced sensitivity to replication-stress-inducing agents than RAD51B-deficient cells, highlighting additive roles of RAD51C in replication stress responses. Overall, these observations confirmed that although FANCM co-depletion rescues DNA damage in RAD51B/RAD51C-deficient cells, cell survival remains unaltered, possibly due to other essential functions of FANCM during replication stress (Collis *et al*, 2008; Ling *et al*, 2016; Lu *et al*, 2019; Luke-Glaser *et al*, 2010; Pan *et al*, 2017; Panday *et al*, 2021; Sahoo *et al*., 2026; Schwab *et al*, 2010).

**Figure 5.**
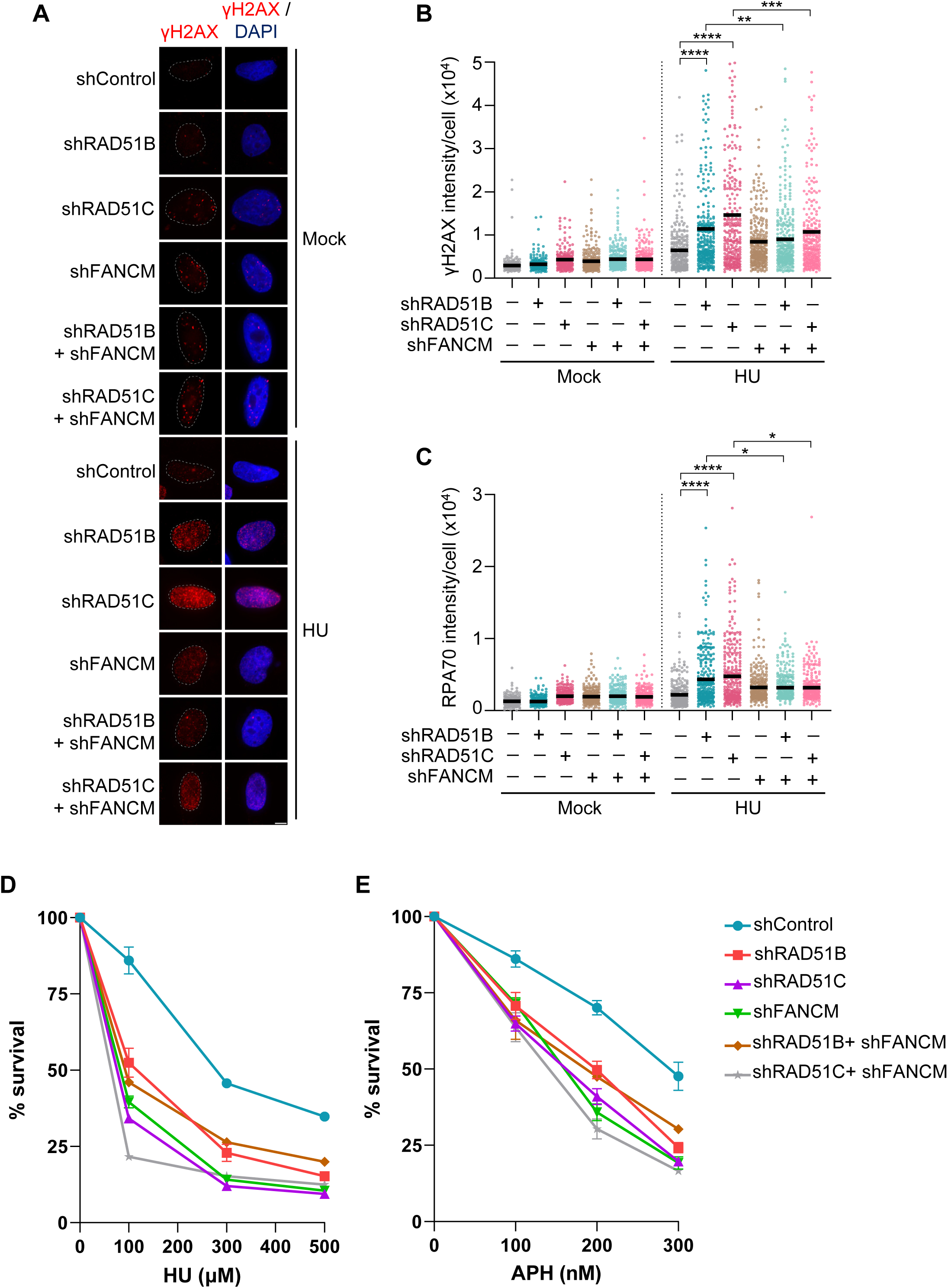
FANCM co-depletion rescues DNA damage in RAD51B/C-deficient cells without rescuing cell survival under replication stress. (A) Representative images showing γH2AX foci formation in the indicated U2OS cells with and without 4 mM HU treatment for 4 h. (B) Quantitative scatter plot of γH2AX intensity per cell in cells as shown in (A). Black bars represent median values. A minimum of 284 cells were counted for each sample from three independent experiments. Mann-Whitney *t* test, **P<0.01, ****P<0.001, ****P<0.0001, P(shControl HU vs shRAD51B HU; shControl HU vs shRAD51C HU) <0.0001, P(shRAD51B HU vs shRAD51B+shFANCM HU) = 0.0082, P(shRAD51C HU vs shRAD51C+shFANCM HU) = 0.0008. (C) Quantitative scatter plot of RPA70 intensity per cell. Black bars represent median values. A minimum of 290 cells were counted from 3 independent experiments. Mann-Whitney *t* test, *P<0.05, ****P<0.0001, P(shControl HU vs shRAD51B HU; shControl HU vs shRAD51C HU) <0.0001, P(shRAD51B HU vs shRAD51B+shFANCM HU) = 0.0231, P(shRAD51C HU vs shRAD51C+shFANCM HU) = 0.0221. (D) Plot showing survival of HeLa cells treated with the indicated shRNAs and increasing doses of HU for 6 days. Cell survival was determined by the MTT assay. (E) Plot showing survival of HeLa cells treated with the indicated shRNAs and increasing doses of APH for 6 days. Cell survival was determined by the MTT assay.

## Discussion

Accurate DNA replication during S-phase is essential for maintaining genome stability. Replication stress response activates multiple fork remodeling and protection mechanisms in the cell, and perturbation in any of these pathways leads to genomic instability and tumorigenesis (Zeman & Cimprich, 2014). Fork stalling is frequently accompanied by fork reversal, a reaction catalyzed by different fork remodeler proteins (Joseph *et al*, 2020). Although reversed forks are physiologically beneficial to cells in dealing with replication stress, these one-ended DSB-like structures require protection from nucleolytic degradation. Numerous DNA damage repair proteins have been shown to protect stalled forks from degradation (Schlacher *et al*., 2011; Schlacher *et al*., 2012; Taglialatela *et al*., 2017; Zellweger *et al*., 2015). The RAD51 paralogs participate in replication stress responses (Saxena *et al*., 2019; Saxena *et al*., 2018; Somyajit *et al*., 2015), yet the mechanisms underlying RAD51 paralog complexes participation in fork protection are unclear. Data presented here demonstrates that distinct complexes of the RAD51 paralogs protect stalled replication forks generated by specific fork remodeling enzymes (Figure 6).

**Figure 6.**
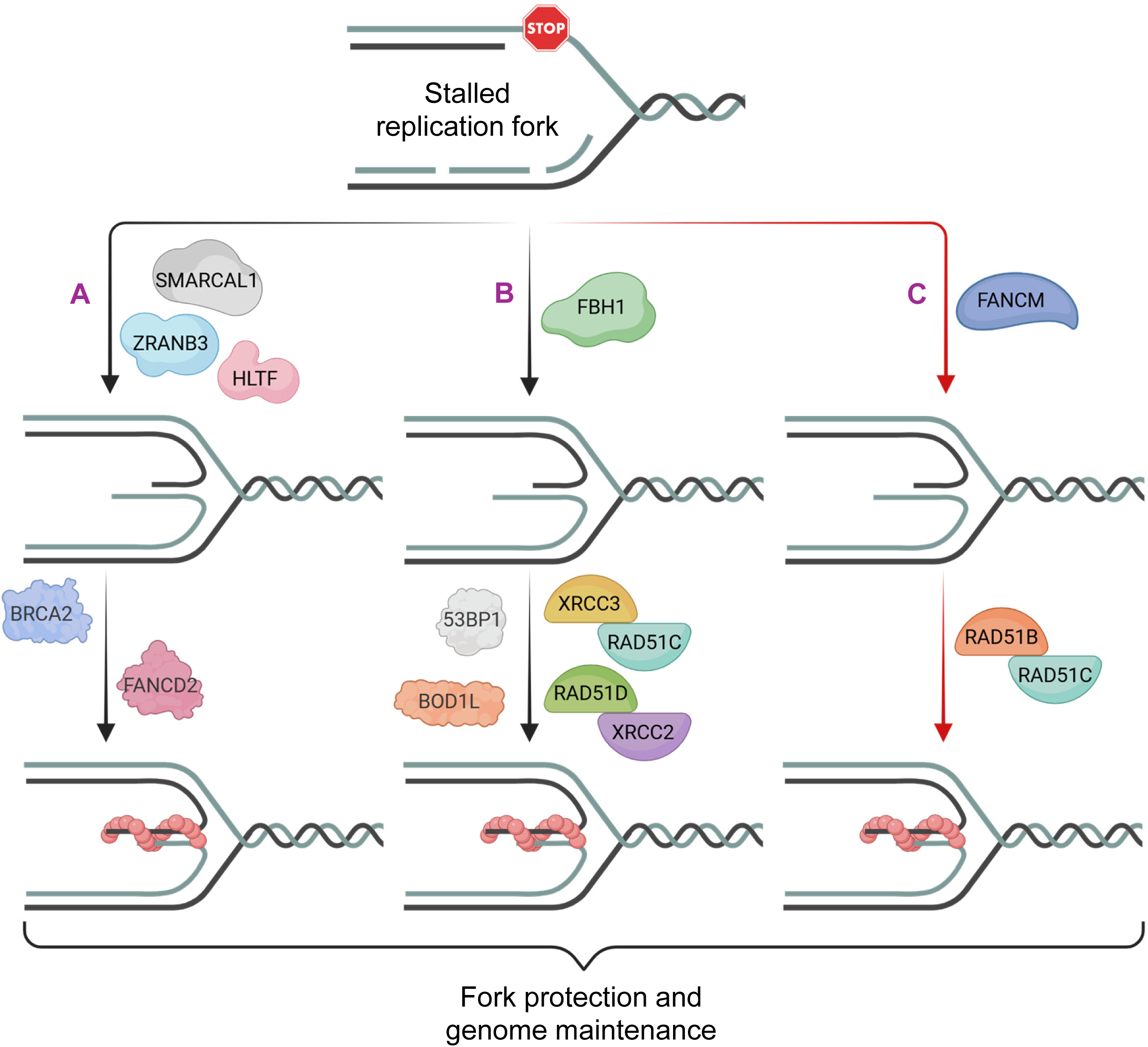
Distinct pathways of stalled fork remodeling and protection during replication stress. Fork reversal can occur via three independent pathways, mediated by the SNF2-family remodelers (A), the FBH1 helicase (B), or the FANCM DNA translocase (C). Each pathway of fork remodeling is accompanied by the recruitment of a distinct set of fork protection factors, which protect reversed fork ends from degradation by nucleases. The RAD51 paralogs participate in the protection of stalled forks in distinct complexes, wherein the DX2 and CX3 complexes protect forks remodeled by the FBH1 pathway (B), and the BC subcomplex protects forks remodeled by the FANCM pathway (C). *This model was created in BioRender*.

Previous studies have established that stalled replication forks remodeled by the SNF2-family members like SMARCAL1, ZRANB3 and HLTF are protected by BRCA1, BRCA2 and FA pathway proteins like FANCD2 and FANCC (Schlacher *et al*., 2011; Schlacher *et al*., 2012; Taglialatela *et al*., 2017). Both the BRCA proteins and the RAD51 paralogs function as mediators of RAD51 nucleofilament formation during HR-mediated DSB repair (Davies *et al*, 2001; Liu *et al*., 2007; Takata *et al*., 2000). In contrast, our data show that BRCA2 and the RAD51 paralogs function in independent pathways of fork protection during replication stress. RAD51 paralog-depleted cells undergo severe fork degradation in cancer-derived U2OS or non-transformed RPE1 cells, and this degradation cannot be rescued by co-depletion of SMARCAL1. This contrasts with BRCA2-depleted cells, where SMARCAL1 co-depletion rescues fork degradation. These observations establish that the RAD51 paralogs protect stalled replication forks independently from the classical BRCA2/SNF2 pathway.

The inability of SMARCAL1 depletion to suppress fork degradation in RAD51 paralog-deficient cells suggested that the paralogs may participate in an alternative pathway of fork protection. Previous studies identified a second fork remodeling pathway mediated by the FBH1 helicase (Liu *et al*., 2020), which is independent of the SNF2-family remodeler pathway. Forks remodeled by FBH1 require factors like 53BP1, BOD1L, VHL and FANCA, rather than BRCA2 or FANCD2, for protection from DNA2-mediated fork degradation (Fugger *et al*, 2015; Liu *et al*., 2020). Surprisingly, FBH1 depletion suppressed fork degradation in RAD51D, XRCC2, and XRCC3-depleted cells, indicating that these paralogs contribute to the protection of FBH1-remodeled forks. In contrast, co-depletion of FBH1 failed to rescue fork degradation in RAD51B-depleted cells. However, RAD51C-depleted cells exhibited a partial but significant rescue with FBH1 co-depletion. RAD51C is a component of multiple paralog complexes, mainly the BCDX2 and CX3 complexes and the BC subcomplex (Braybrooke *et al*, 2000; Chun *et al*., 2013; Masson *et al*., 2001b; Miller *et al*., 2002; Sigurdsson *et al*., 2001; Takata *et al*., 2000; Wiese *et al*, 2002). Notably, distinct complexes of RAD51 paralog have been shown to perform specialized functions during replication stress, with the DX2 subcomplex regulating replication fork progression under conditions of limited nucleotide availability (Saxena *et al*., 2018), and the CX3 complex playing a key role in the restart of the stalled forks (Berti *et al*, 2020b; Petermann *et al*, 2010; Somyajit *et al*., 2015), regulation of replication speed under DNA-damaging treatment (Henry-Mowatt *et al*, 2003), and resolution of R-loops (Sahoo *et al*., 2026). Our data establish that the DX2 and CX3 sub-complexes protect forks remodeled by the FBH1 helicase. These observations support a functional specialization of distinct RAD51 paralog complexes during replication stress responses. Recent cryo-electron microscopy-based studies have identified additional RAD51 paralog assemblies (Greenhough *et al*, 2025; Koo *et al*, 2026; Rawal *et al*, 2026). Greenhough et al. identified an X3CDX2 complex and showed that this complex stabilizes the 5′ termini of RAD51 nucleofilaments, while the BCDX2 helps in ATP-dependent RAD51 filament formation (Greenhough *et al*., 2025). Another study identified a BCDX2-CX3-RAD51 super-complex and characterized it as a protofilament template for RAD51 filament assembly (Koo *et al*., 2026). However, whether CX3 and DX2, as subcomplexes or as the X3CDX2 complex, contribute to the protection of FBH1-remodeled forks remains to be investigated. Interestingly, ATR-mediated phosphorylation of XRCC2 and XRCC3 differentially regulates the role of the RAD51 paralog complexes in replication fork progression and DSB repair, respectively (Saxena *et al*., 2019; Saxena *et al*., 2018; Somyajit *et al*., 2013). Whether ATR-mediated phosphorylation regulates the assembly of distinct complexes of RAD51 paralogs and their function in stalled fork protection remains to be investigated.

A striking observation of our study is that BC subcomplex protects stalled forks remodeled by the FANCM DNA translocase. FANCM is a bona fide FA pathway protein that facilitates the assembly of the FA core complex at interstrand cross-link (ICL) sites and subsequent monoubiquitination of FANCD2 during ICL repair (Meetei *et al*, 2005) (Ciccia *et al*, 2007). Interestingly, FANCM-deficient patient cells were found to be sensitive to a wide range of genotoxic agents, such as camptothecin (CPT) and UV, despite retaining FANCD2 monoubiquitination (Singh *et al*, 2013), suggesting that FANCM contributes to genome stability through mechanisms distinct from the canonical FA pathway. FANCM has ATP-dependent bidirectional DNA translocase activity and binds branched DNA structures to remodel them into four-way junctions in vitro (Gari *et al*., 2008; Xue *et al*, 2008; Yan *et al*, 2010). Deficiency of FANCM has been linked to unregulated fork progression under both unperturbed conditions and low levels of replication stress (Luke-Glaser *et al*., 2010), underscoring the importance of FANCM’s role in regulating replication dynamics. Physical barriers, such as UV lesions or ICLs, generate torsional stress ahead of the replication fork, and fork reversal mediated by FANCM may help alleviate this strain while providing access for DNA repair enzymes to damaged template strands (Luke-Glaser *et al*., 2010). Consistent with this model, FANCM has also been implicated in the efficient restart of stalled replication forks (Schwab *et al*., 2010), raising the possibility that its bidirectional translocase activity may facilitate both fork reversal and subsequent fork restart depending on the cellular context. FANCM is recruited to ICL-induced stalled forks through its interaction with the replisome component AND1 (Zhang *et al*, 2022), indicating that ICL-induced fork stalling triggers FANCM-mediated fork reversal. Our results show that stalled forks reversed by FANCM specifically require the BC subcomplex for protection, whereas BRCA2 or 53BP1/BOD1L do not participate in this pathway in U2OS cells. These observations support the existence of multiple mechanistically distinct fork remodeling pathways in cells that recruit specialized fork protection factors.

Interestingly, we find complete rescue of fork degradation in RAD51C-deficient cells with FANCM co-depletion, but only partial rescue with FBH1 co-depletion. A possible explanation is that RAD51C predominantly participates in fork protection through the BC complex and, to a lesser extent, through the CX3 complex. Partial suppression of fork degradation upon FANCM co-depletion in BRCA2-deficient RPE1 cells, but not in U2OS cells, imply a possible cell-type dependence of these pathways. Similar cell-lineage-specific differences have been reported for 53BP1, where it has been shown to be important for fork protection in U2OS but not in RPE1 cells (Liu *et al*., 2020). Interestingly, a recent study reported profound synthetic lethality between SMARCAL1 and FANCM, as combined depletion of both proteins leads to severe genomic instability (Feng *et al*, 2024). Considering our findings, it becomes plausible that the complementary functions of distinct fork remodeling pathways give rise to such synthetic lethality, where FANCM-mediated fork reversal becomes particularly crucial for maintaining genome stability under replication stress in the absence of SMARCAL1 or vice versa. How specific fork protection factors are recruited to stalled forks in a remodeler-specific manner remains an important open question, and further studies examining the structural and genomic contexts of these remodeling events can provide further insight into the mechanisms of fork protection during replication stress.

RAD51 is essential for fork reversal in response to genotoxic stress (Zellweger *et al*., 2015). Recent studies have shown that, through its strand-exchange activity, RAD51 bypasses the CMG helicase to form a parental strand duplex, which is then acted upon by fork remodelers to generate reversed fork structures (Liu *et al*., 2023). Consistent with the upstream role of RAD51 in fork reversal, RAD51 depletion rescued the fork degradation in RAD51B- and RAD51C-depleted cells, indicating that FANCM-mediated fork reversal is dependent on RAD51 activity. In BRCA2-deficient cells, the absence of stable RAD51 nucleofilaments at the reversed fork ends leads to pathological fork degradation by MRE11 and EXO1 (Lemacon *et al*., 2017). Additionally, the DNA2 endonuclease has been shown to resect reversed DNA ends to promote fork restart, independently of the MRE11 pathway (Thangavel *et al*., 2015). Liu et al. demonstrated that BRCA2-deficient cells can undergo fork degradation by both MRE11 and DNA2 nucleases, but FBH1-remodeled forks are preferentially degraded by DNA2 (Liu *et al*., 2020). In our experiments, fork degradation in RAD51B/C-depleted cells was rescued by treatment with mirin, C5, or shEXO1, suggesting that RAD51B/C protects stalled forks from nucleolytic attack by multiple nucleases. Whether DNA2-mediated fork processing is essential for restarting stalled forks in the FANCM-mediated pathway remains to be investigated.

In the absence of proper fork protection or fork restart, stalled forks can accumulate ssDNA stretches or collapse into DSBs (Hanada *et al*., 2007; Kolinjivadi *et al*, 2017b; Pasero & Vindigni, 2017; Zellweger *et al*., 2015). Our studies showed that RAD51B/C-deficient cells accumulate high amounts of DNA damage markers, which were rescued upon FANCM co-depletion, indicating that FANCM-mediated fork remodeling causes DNA damage in these cells. Interestingly, although FANCM co-depletion suppresses the accumulation of DNA damage in RAD51B/C-deficient cells, it does not restore cellular survival following replication stress. This separation between DNA damage suppression and cell viability suggests that FANCM performs additional essential functions in the replication stress response beyond fork remodeling. Indeed, FANCM has been implicated in multiple genome maintenance processes, including coordination of replication restart, resolution of TRCs, and recruitment of FA pathway components to stalled forks and fork repair (Blackford *et al*, 2012; Luke-Glaser *et al*., 2010; Panday *et al*., 2021; Sahoo *et al*., 2026; Schwab *et al*., 2010; Xue *et al*, 2015). Together, these observations indicate that the critical roles of FANCM and the RAD51 paralogs remain indispensable for cell survival under replication stress.

Collectively, our findings reveal an unexpected functional specialization among the RAD51 paralog complexes in the protection of stalled replication forks. Distinct RAD51 paralog subcomplexes operate in remodeler-specific fork protection pathways, with the DX2 and CX3 (or the X3CDX2) complexes protecting FBH1-remodeled forks, while the BC subcomplex safeguards forks remodeled by FANCM (Figure 6). These observations support a model in which the identity of the fork remodeler dictates the recruitment of specialized fork protection factors. Conceivably, cells have evolved diverse fork-protection mechanisms to tackle a wide range of replication impediments encountered during genome duplication. However, several important questions remain unresolved. It is unclear how distinct fork remodelers generate topologically different fork intermediates that selectively recruit specific protection factors. How the different fork remodeling pathways cooperate spatially and temporally remain to be investigated. Furthermore, the molecular mechanisms through which individual RAD51 paralog complexes stabilize these remodeled forks remain to be elucidated, including whether they act through modulation of RAD51 filament dynamics alone or if they directly regulate nuclease access to nascent DNA. Addressing these questions will be critical for understanding how multiple fork protection pathways cooperate to preserve genome stability during replication stress.

## Methods

### Cell lines and culture conditions

Human cell lines U2OS, HeLa and RPE1 were grown in Dulbecco’s Modified Eagle Medium (DMEM) supplemented with 10% FBS, 1% penicillin/streptomycin (Sigma-Aldrich) and 1% Glutamax (Gibco) at 37°C in a humidified air chamber containing 5% CO2.

### Plasmids and transfections

All plasmid transfections for transient depletion/expression were performed using a Bio-Rad gene pulsar X cell (260 V and 1050 μF). Fresh media was added to the cells 6-8 h after transfection. Cells were processed for indicated treatments/experiments 24-30 h after transfection. All shRNAs were generated using previously reported sequences (Table S1) and cloned into the pRS shRNA vector. The RAD51 mutants used were reported previously (Dixit *et al*, 2024; Nagraj *et al*, 2025).

### DNA fiber assay

The DNA fiber assay was performed as previously described (Bhattacharya & Nagaraju, 2026; Dixit *et al*., 2024; Nagraj *et al*., 2025). Briefly, cells were sequentially pulse-labeled with 25 µM CldU (Sigma) and 250 µM IdU (Sigma) followed by 4 mM HU treatment for 5 h for fork protection assay. Next, the cells were incubated in ice-cold PBS for 10 mins, harvested, counted and re-suspended in 250 μl PBS. 3 ul of the cell mixture was mixed with 7 µl of lysis buffer on glass slides (ThermoScientific superfrost) and allowed to stand for 7 min. Slides were inclined at 45° to spread the cell suspension, and the slides were fixed in a methanol:acetic acid (3:1) solution at 4°C, O/N. The following day, DNA was denatured by incubating in 2.5 M HCl for 1 h and blocked with 2% BSA in 0.1% PBST solution (1X PBS and 0.1% Tween-20). Next, the slides were incubated with primary antibodies for 2.5 h and secondary antibodies for 1 h at RT. Coverslips were mounted on the slides with a Mowiol mounting medium (Sigma) and visualized using an Apotome microscope (Zeiss Axio observer). Antibodies used for performing DNA fiber studies include: rat anti-BrdU for CldU (1:500, Abcam), mouse anti-BrdU for IdU (1:250, BD Biosciences), rabbit anti-mouse IgG (Alexa Fluor 488) (1:500, Abcam) and donkey anti-rat IgG (Alexa Fluor 594) (1:500, Abcam).

### Immunofluorescence

Exponentially growing cells were seeded onto coverslips 8 h after transfection, and treated as indicated. Next, cells were washed with PBS, pre-extracted with 0.5% Triton X-100 for 90 s on ice and fixed in 4% formaldehyde for 10 min at RT. After three PBS washes, coverslips were blocked in blocking buffer (0.5% BSA and 0.5% Triton X-100 in PBS) for 30 min. The coverslips were incubated with the indicated primary antibodies for 2 h at RT. After a wash with blocking buffer, the coverslips were incubated with respective FITC/TRITC-conjugated secondary antibodies for 1 h at RT and then stained with DAPI (1 µg/ml; Sigma-Aldrich) for 10 min before mounting onto slides with Mowiol 4-88 (Sigma). Images were acquired using an Apotome microscope (Zeiss Axio observer) and processed using ImageJ software. The following antibodies were used for performing immunofluorescence experiments in this study: rabbit anti-RPA70 (1:2000, Abcam) and mouse anti-H2AX-pS139 (1:2000, BD Biosciences).

### Cell survival assay

Cells were plated at 3000 cells/well in 96-well plates 12 h after shRNA transfection. Appropriate doses of HU and APH were added 24 h post-transfection, and cells were continuously treated for 5-7 days. Later, cell survival was measured by MTT (0.25 mg/ml; Sigma-Aldrich) assay using a microplate reader (VersaMaxROM version 3.13). Percent cell survival was calculated as treated cells/untreated cells*100.

### Western blotting

Immunoblotting was performed as previously described (Nath & Nagaraju, 2020). Cells were harvested and lysed in RIPA lysis buffer (50 mM Tris-HCl pH 7.5, 1% NP40, 0.5% sodium deoxycholate, 0.1% SDS, 150 mM NaCl, 2 mM EDTA, and 50 mM sodium fluoride) supplemented with cOmplete mini protease inhibitor cocktail (Roche). Protein quantity was estimated by standard Bradford assay. 50 μg of protein samples were resolved on SDS-PAGE gel and transferred onto PVDF membranes (Millipore) by semi-dry transfer method (Bio-Rad Trans-Blot SD). Membranes were blocked using 5% skim-milk (Hi-media) in TBST (50 mM Tris-HCl, pH 8.0, 150mM NaCl, 0.1% Tween 20) and incubated with primary antibody overnight (O/N) at 4°C or 2-3 h at RT, followed by HRP-conjugated secondary antibody incubation for 1 h at room temperature (RT). After three TBST washes, membranes were developed with chemiluminescent HRP substrate (Millipore) and imaged using Chemidoc (Bio-Rad Chemidoc Imaging System). The following primary antibodies were used in this study for western blotting: mouse anti-RAD51C (1:250, SC), mouse anti-RAD51D (1:200, SC), mouse anti-RAD51B (1:200, SC), mouse anti-XRCC2 (1:250, SC), mouse anti-XRCC3 (1:200, SC), rabbit anti-FANCM (1:2000, Abcam; 1:500, Invitrogen), mouse anti-FBH1 (1:500, SC), rabbit anti-SMARCAL1 (1:500, Abcam), rabbit anti-ZRANB3 (1:500, Abcam), mouse anti-HLTF (1:500, SC), rabbit anti-RAD51 (1:500, Abcam), rabbit anti-BRCA2 (1:500, Abcam), rabbit anti-53BP1 (1:500, novus), mouse anti-Flag (1:1000, Sigma), rabbit anti-EXO1 (1:500, Abcam), and mouse anti-MCM3 (1:2000, SC).

## Acknowledgements

We thank members of the GN lab for their useful discussions. We thank Kumar Somyajit for his suggestions on the manuscript. Funding by Department of Science and Technology (DST) (CRG/2022/003533); Department of Atomic Energy (58/14/03/2022-BRNS); Department of Biotechnology (DBT) (BT/PR45508/MED/30/2414/2022); Council of Scientific and Industrial Research (CSIR) (37/1756/23/EMR-II); J.C. Bose fellowship (JCB/2021/000009), Indian Institute of Science (IISc)-DBT partnership program (BT/PR27952/INF/22/212/2018) and infrastructure support provided by funding from DST and UGC are greatly acknowledged. D.B. was supported by a fellowship from DST, IISc and J. C. Bose Fellowship. H.K.D. was supported by a fellowship from IISc. S.S was supported by a fellowship from CSIR, IISc and DBT. S.K. was supported by a fellowship from IISc.

## Author contributions

D.B. and G.N. conceived the project and designed the experiments. D.B., H.K.D., S.S. and S.K. performed the experiments. D.B., H.K.D., S.S. and G.N. analyzed the data. D.B. wrote the original draft of the manuscript. G.N. reviewed and edited the manuscript.

